# CXCR4-associated depletion of bone marrow CD34^+^ cells following CCR5-tropic HIV-1 infection of humanized NOD/SCID/JAK3^null^ mice and partial protection of those cells by a promotor-targeting shRNA

**DOI:** 10.1101/190702

**Authors:** Tetsuo Tsukamoto

## Abstract

**Objectives:** Hematological abnormalities that include changes in bone marrow, such as in anemia and pancytopenia, are common among human immunodeficiency virus (HIV)-infected patients, particularly in the advanced stage of disease. This study aimed to provide better experimental evidence of such manifestations in animal models.

**Design:** NOD/SCID/JAK3^null^ (NOJ) mice were transplanted with human cord-derived CD34^+^ cells with or without transduction with a lentiviral vector expressing a promoter-targeting shRNA called PromA.

**Methods:** At 16 weeks after transplantation, mice engrafted with CD34^+^ cells were infected with CCR5-tropic HIV-1_JRFL_.

**Results:** At week 2 post infection, HIV replication was observed in peripheral blood mononuclear cells and splenocytes. In mice transplanted with unmanipulated CD34^+^ cells, viral replication was accompanied by a loss of peripheral/spleen CD4^+^CCR5^+^ T cells. Interestingly, bone marrow CD34^+^ cells in HIV-infected mice were also depleted, but in a CXCR4-associated manner. Conversely, the lentiviral transfer of PromA in CD34^+^ cells prior to transplantation rendered the humanized NOJ mice resistant to HIV replication in CD4^+^ T cells, resulting in better preservation of peripheral/spleen CD4^+^CCR5^+^ T cells and bone marrow CD34^+^ cells at two weeks after infection.

**Conclusions:** These results implicate the importance of evaluating hematopoietic stem/progenitor cell pools in addition to peripheral CD4^+^ T-cell counts to assess the early stage of HIV infection. Moreover, stable gene transfer of PromA to hematopoietic stem cells not only limited HIV replication but also led to preservation of different subsets of hematopoietic cells, including bone marrow stem/progenitor cells.

## Introduction

Human immunodeficiency virus type 1 (HIV-1) infection induces hematological changes [1]. Although antiretroviral therapy (ART) is effective for viremia control and acquired immunodeficiency syndrome (AIDS) treatment, some patients fail to achieve sufficient restoration of T-cell immune function despite being aviremic during treatment [2]. Such immunological nonresponsiveness has been associated with immune activation and bone marrow impairment [3, 4]. Bone marrow abnormalities, such as dysplasia and abnormal hematopoietic cell development, are frequently observed in HIV-infected individuals [5]. Therefore, it is desirable to design a method to protect the whole hematopoietic cell population from HIV infection-associated immunological and hematological disorders.

Transcriptional gene silencing (TGS) occurs through small non-coding RNAs that direct epigenetic changes, such as DNA methylation and heterochromatin formation. TGS was first described in plants [6] and has since been adapted in humans for the development of new therapeutic agents [7]. For example, previous studies have reported sustained, profound, and highly specific suppression of viral replication by small interfering RNA (siRNA)- and short hairpin RNA (shRNA)-induced TGS of HIV-1 and simian immunodeficiency virus in various *in vitro* models via a mechanism that results in chromatin compaction [8-14]. HIV-1 has identical long terminal repeats (LTRs) at the 5’ and 3’ ends of the integrated virus; therefore, any promoter-targeted siRNA can potentially act through post-transcriptional gene silencing (PTGS). However, the use of a lead candidate, PromA, showed that the contribution of PTGS is limited [8].

In a previous collaborative study including *in vivo* animal experiments, a lentiviral delivery system was used to express the previously described shRNA PromA targeting the HIV-1 promoter region to transduce human peripheral blood mononuclear cells (PBMCs), whereby an antiviral effect of TGS produced by this construct on HIV-1 infection *in vivo* was demonstrated using NOD/SCID/Jak3^null^ (NOJ) mice [15] transplanted with the lentivirally transduced PBMCs [13]. In the present study, we tested whether TGS using shRNA (shPromA) can be applied to more profound gene therapies, such as gene-modified hematopoietic stem cell transplantation to replace infected immune cells with HIV-resistant cells. Humanized mouse models have been characterized for tissue distribution and HIV infection of human hematopoietic cells [16]. To better utilize the NOJ-based humanized mouse model, irradiated newborn NOJ mice were transplanted with human cord-derived CD34^+^ cells, as previously described [15]. It is demonstrated in the present study that the generated hematopoietic/immune cells, particularly CD4^+^ T cells, were resistant to replication of the challenged HIV-1_JRFL_ *in vivo* and that bone marrow CD34^+^ cells expressing shPromA may be better protected and preserved after HIV infection.

## Materials and Methods

### Virus stocks

HIV-1_JRFL_ stocks were produced by 293T-cell transfection with the molecular DNA clone pJRFL. After transfection, the culture supernatant was collected and viral titers were determined using an HIV p24 Enzyme-Linked ImmunoSorbent Assay (ELISA) kit (Zeptometrix Corporation, Buffalo NY, USA).

### Cells

Umbilical cord blood samples were collected from healthy newborn infants at Fukuda Hospital (Kumamoto, Japan) after obtaining informed consent. Cord blood mononuclear cells were isolated using Pancoll separation solution (PAN-Biotech GmbH, Aidenbach, Germany) and density gradient centrifugation. Labelled CD34^+^ cells were isolated using human CD34 micro beads and LS columns (Miltenyi Biotec, Macquarie Park, NSW, Australia). The purity of the CD34^+^ cells was consistently >92% by flow cytometric analysis.

### Lentiviral production

Self-inactivating lentiviral vectors expressing shRNA PromA (shPromA) or PromA M2 (shPromA-M2) (Fig. 1) were constructed, as previously described [13]. Vesicular stomatitis virus-G pseudotyped lentiviral vectors were prepared by plasmid DNA transduction into 293T cells using Lipofectamin 2000 reagent (Thermo Fisher Scientific, Scoresby, VIC, Australia). The viruses were concentrated from supernatant, as previously described [17, 18], and stocks were titrated on 293T cells based on green fluorescent protein (GFP) expression.

**Fig 1.**
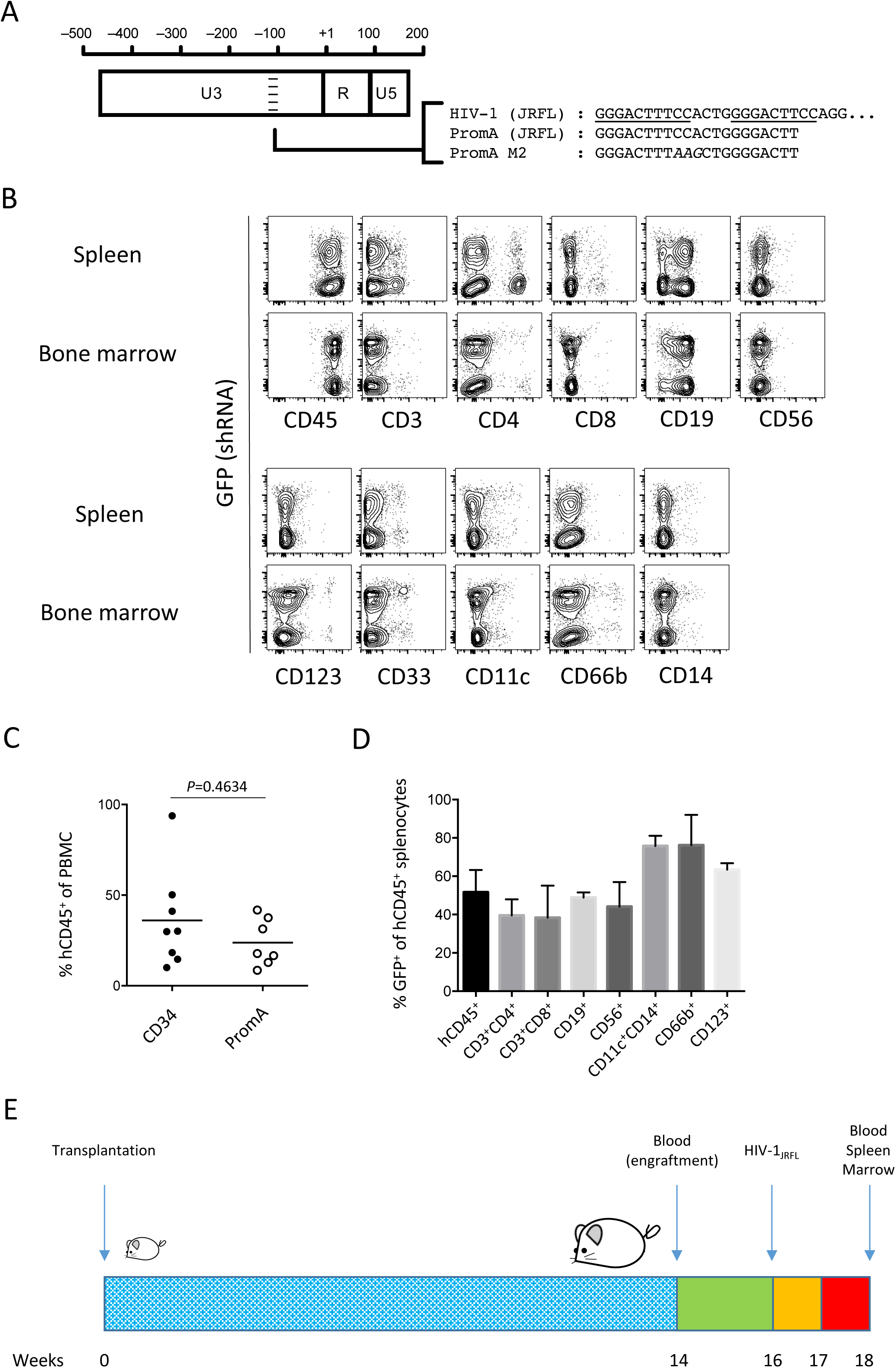
Design and plan of the study. (A) A schematic representation of the siRNAs PromA (JRFL) and PromA-M2. The two NF-kB binding sites in the HIV-1_JRFL_ proviral DNA sequence are underlined. The three mismatched bases in PromA-M2 compared with PromA_JRFL_ are indicated with italic letters. (B) Three mice were sacrificed without HIV challenge 15 weeks after transplantation with shPromA-transduced cells. Splenocytes and bone marrow cells were collected from three mice, and different major subsets of human CD45^+^ hematopoietic cells were screened for GFP expression. Representative plots showing both GFP^+^ and GFP^-^ cells in all of the subsets of human hematopoietic cells tested. (C) Comparison of human CD45^+^ frequencies between the CD34 group (transplanted with unmanipulated CD34^+^ cells) and the PromA group (transplanted with shPromA-transduced cells) mice that were included in the following HIV challenge experiment. (D) The mean shRNA-expressing (GFP^+^) percentages in the different subsets of cells in the spleen. Statistical analysis was performed with the Mann–Whitney U-test. (E) The *in vivo* study plan. Newborn NOJ mice were irradiated and transplanted with PromA-transudced or unmanipulated CD34^+^ cells. At 14 weeks after treatment, blood was collected from the retro-orbital sinus and the PBMCs were tested for human CD45 expression. Two weeks later, mice were intraperitoneally challenged with HIV-1_JRFL_. Blood was collected at weeks 1 and 2 post challenge. Mice were sacrificed at week 2 for the collection of splenocytes and bone marrow cells.

### CD34^+^ cell transduction with lentiviruses

After culturing in X-VIVO 10 media (Lonza, Sydney, NSW, Australia), fresh cord-derived CD34^+^ cells were transduced with a lentiviral vector at a multiplicity of infection (MOI) of 10 following pre-stimulation for 24 h with 50 ng/mL each of stem cell factor (SCF), FMS-related tyrosine kinase 3 ligand (Flt-3L), and thrombopoietin (TPO) (R&D Systems, Inc., Minneapolis, MN, USA). The cells were then collected and cryopreserved. A small portion of cells was further incubated for 48 h to check the transduction rates by measuring GFP expression levels by flow cytometry.

### Cord-derived human CD34^+^ cell transplantation into NOJ mice and HIV-1 infection

Humanized NOJ mice were generated, as previously described [15]. Briefly, newborn NOJ mice were irradiated (1.0 Gy) and then intrahepatically infused with shPromA-transduced or unmanipulated CD34^+^ cells (2 × 10^5^) that were resuspended in phosphate-buffered saline (0.1 mL). At week 14 after transplantation, peripheral blood was collected from the retro-orbital sinus for flow cytometric analysis of PBMCs to check the engraftment of human hematopoietic cells (Fig. 3). At week 16 after transplantation, each humanized mouse was intraperitoneally inoculated with 200 ng of HIV-1_JRFL_ p24, as determined with the HIV-1 p24 antigen ELISA (Fig. 3). Blood was collected at weeks 1 and 2 after infection to obtain plasma and PBMCs. At week 2, the mice were sacrificed, and the splenocytes and bone marrow cells were collected from the left femur. All animal experiments were performed in accordance with the guidelines of the Kumamoto University Graduate School of Medical Science.

**Fig 3.**
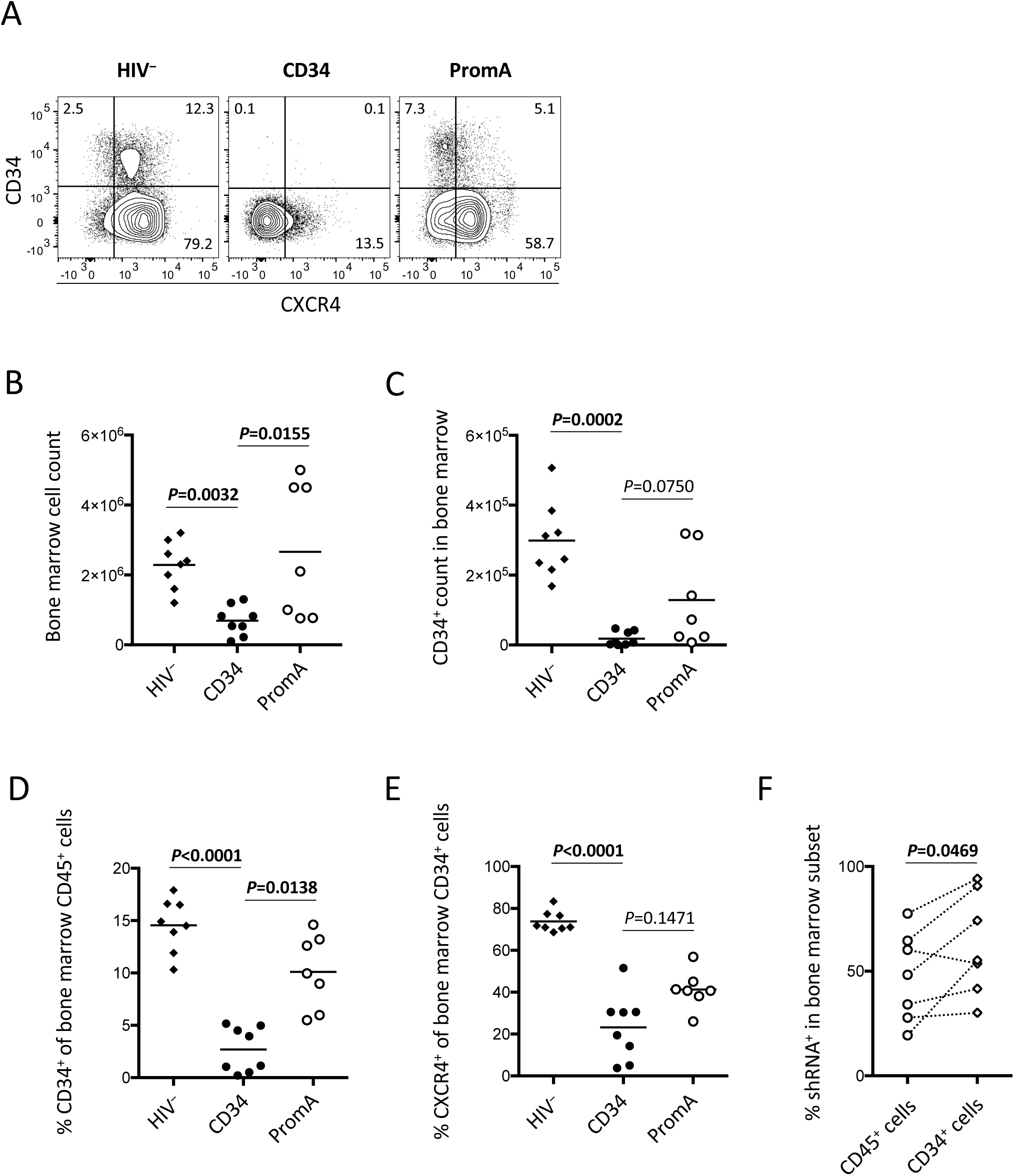
Bone marrow cell counts and percentages in humanized mice. Cells were collected from the left femur at week 18 post transplantation for uninfected mice or at week 2 post infection, counted, and analyzed by flow cytometry. (A) Representative plots showing the CD34/CXCR4 expression patterns in untransduced uninfected (left), untransduced infected (middle), and shPromA-transduced infected (right) mice. (B) Bone marrow cell counts. (C) Bone marrow CD34^+^ cell counts. (D) Percentages of CD34^+^ cells in the population of bone marrow CD45^+^ cells. (E) Percentages of CXCR4^+^ cells in the population of bone marrow CD34^+^ cells. (F) Comparison of shPromA^+^ frequencies in CD45^+^ and CD34^+^ bone marrow cells of mice in the PromA group. Statistical analyses were performed by the nonparametric multiple comparison analysis with Dunn’s method (B-E) or the Wilcoxon matched-pairs signed rank test (F).

### PCR/RT-PCR analysis

Cellular DNA was extracted using the Kaneka Easy DNA Extraction Kit (Kaneka Corporation, Osaka, Japan). Cellular RNA was extracted using the High Pure RNA Tissue Kit (Roche Diagnostics, Tokyo, Japan). Extracted DNA and RNA were amplified by PCR/RT-PCR using an HIV *gag* primer set (sense: 5’-AGTGGGGGGACATCAAGCAGCCATGCAAAT-3’, antisense: 5’-TACTAGTAGTTCCTGCTATGTCACTTCC-3’), as previously described [9]. Both HIV DNA and RNA levels were normalized against glyceraldehyde 3-phosphate dehydrogenase (GAPDH). All quantitative PCR/RT-PCR analyses were performed using a syber green method with an ABI PRISM Genetic Analyzer (Thermo Fisher Scientific).

### Flow cytometric analysis of CD4^+^/CD8^+^ T cells and intracellular p24-expressing cells

Blood samples of HIV-infected NOJ mice were treated with red cell lysing buffer (155 mM NH_4_Cl, 10 mM KHCO_3_, and 0.1 mM EDTA) for 15 min and then resuspended in staining media containing 2% fetal bovine serum (FBS). Cells were stained with Live/Dead Fixable Near-IR Dead Cell Stain dye (Thermo Fisher Scientific) and monoclonal antibodies against mouse CD45 and human CD45, CD3, CD4, CD8, and CCR5 for 15 min and then analyzed using a FACS LSR II Flow Cytometer (BD Biosciences, San Jose, CA, USA). The flow cytometry data were further analyzed using FlowJo v10.8 software (FlowJo, LLC, Ashland, OR, USA).

### Sequence analysis

The 5’ LTR region of the viral RNA was amplified by PCR using a viral RNA-specific primer set (sense: 5’-GACCATCAAGCGGCTATGCAGATT-3’, antisense: 5’-GTAAATGTTGCCTACTGGTATGGG-3’), followed by sequence analysis of the region around the two NF-κB binding sites using the BigDye™ Terminator v3.1 Cycle Sequencing Kit and a sequence analyzer (Thermo Fisher Scientific).

### *In vitro* generation and HIV-1_JRFL_ infection of macrophages

After transduction with either shPromA or shPromA-M2, CD34^+^ cells were stimulated and expanded for two or three weeks in 12-well plates in the same X-VIVO 10 media supplemented with 4% FBS, 50 ng/mL SCF, 15 ng/mL TPO, 30 ng/mL Flt-3L, and 30 ng/mL IL-3 [19]. The cells were then washed and further cultured for 4–7 days in Roswell Park Memorial Institute 1640 media supplemented with 20% FBS, 25 ng/mL SCF, 30 ng/mL Flt-3L, 30 ng/mL IL-3, and 30 ng/mL macrophage colony-stimulating factor. Development of shPromA-expressing macrophages was detected under a fluorescence microscope as GFP^+^ cells with typical fried-egg shapes (Fig. 6A). Cells were then seeded at 1 × 10^5^ per well in a 96-well plate and infected with 50 ng (p24) of HIV-1_JRFL_. Cells were collected five days after infection, stained for surface markers and intracellular HIV p24, and analyzed by flow cytometry.

### Antibodies and reagents for flow cytometry

Anti-human CD34 APC and anti-human CD38 PE (BD Biosciences) were used for the analysis of magnetically separated CD34^+^ cells. The following antibodies were used for the flow cytometric analysis of NOJ mouse PBMCs, splenocytes, and bone marrow cells after human CD34^+^ cell transplantation: anti-mouse CD45 PerCP-Cy5.5, anti-mouse CD45 PE, anti-human CD45 Brilliant Violet (BV) 510, anti-human CCR5 PE-CF594, anti-human CD56 PerCP-Cy5.5, anti-human CD11c BV510, anti-human CD19 PE-CF594, anti-human CD33 PE-Cy7, anti-human CD66b Alexa Fluor 647 (AF647) (BD Biosciences), anti-human CD3 PE-Cy7, anti-human CD4 BV421, anti-human CD8 AF647, anti-human CD14 PerCP-Cy5.5 (BioLegend, San Diego, CA, USA), anti-human CD123 PE (Thermo Fisher Scientific), and anti-HIV-1 p24 PE (Beckman Coulter, Tokyo, Japan). In addition, the following antibodies were used to stain *in vitro*-generated macrophages: anti-human CD4 APC, anti-human CD11b BV421, anti-HLA-DR BV510 (BD Biosciences), anti-human CD14 PE-Cy7, anti-human CD16 AF700, and anti-human CD163 PerCP-Cy5.5 (BioLegend). Dead cells stained with the Live/Dead Fixable Near-IR Dead Cell Stain Kit (Thermo Fisher Scientific) were excluded from analysis.

### Statistical analysis

Statistical analysis was performed using GraphPad Prism software version 7 (GraphPad Software, Inc., La Jolla, CA, USA). A probability (*p*) value of <0.05 was considered statistically significant. The unpaired Mann–Whitney test or Wilcoxon matched-pairs signed rank test was used for comparisons between two samples. Nonparametric multiple comparison analysis was performed using Dunn’s method.

## Results

### Cord-derived CD34^+^ cell transduction with lentiviruses expressing shPromA customized for HIV-1_JRFL_

The siRNAs PromA and PromA M2 are described previously [13]. There was a one-base mismatch in the PromA region between HIV-1_NL4-3_ and HIV-1_JRFL_; therefore, the present study utilized a version of PromA customized for HIV-1_JRFL_, which is simply referred to as PromA in the following text. The sequences of HIV-1_JRFL_, PromA, and PromA-M2 are described in Fig. 1A. The lentiviral vectors expressing PromA and PromA M2 in the shRNA forms (shPromA and shPromA-M2, respectively) were prepared for CD34^+^ cell transduction.

The volumes of the collected cord blood samples ranged from 20 to 130 mL. Each sample contained approximately 5 × 10^6^–2 × 10^8^ mononuclear cells with CD34^+^ cells accounting for about 0.1%–6%, as determined by flow cytometric analysis. After mononuclear cell isolation and CD34^+^ microbead selection, approximately 5 × 10^4^–2 × 10^6^ CD34^+^ cells were obtained. The average purity of the CD34^+^ cells after selection was 96.7%, whereas the average CD38^low^ frequency was 25.6% (Fig. S1A and C). The CD34^+^ cell batches with >4 × 10^5^ cells were used for lentiviral shPromA transduction. Fresh CD34^+^ cells were pre-stimulated in X-VIVO 10 media containing SCF, Flt-3L, and TPO for 24 h and then transduced with lentiviruses expressing shPromA at an MOI of 10. Cells were cryopreserved 24 h after transduction. A small portion of cells was cultured for another 48 h to evaluate transduction efficiency, which was expressed as the percentage of GFP^+^ cells in the culture. The average GFP^+^ frequency was 65.1% (Fig. S1B and C). The PromA-transduced CD34^+^ cells exhibited relevant *in vitro* colony formation capacity compared with untransduced CD34^+^ cells (Fig S2).

### Engraftment of NOJ mice with shPromA-expressing cells

Newborn NOJ mice were irradiated and intrahepatically transplanted with CD34^+^ cells. At 12–14 weeks after transplantation, peripheral blood was obtained from the retro-orbital sinus. PBMCs were isolated and analyzed by flow cytometry. Twenty mice transplanted with shPromA-transduced cells (PromA group) and 12 transplanted with unmanipulated CD34^+^ cells (CD34 group) were alive during engraftment assessment. After excluding poorly engrafted mice with low human CD45^+^ frequencies (<3.0%) or poorly differentiated CD3^+^ cells, the percentages of human CD45^+^ cells were compared between the PromA and CD34 groups, but no significant difference was detected (Fig. 1C). These mice were included in the following HIV challenge experiment (Fig. 1E). The average GFP^+^ frequency of human CD45^+^ cells was 40.8% in the PromA group (Fig. S3A), which was lower than the average transduction efficiency measured prior to transplantation (65.1%, Fig. S1C). Three of the mice transplanted with shPromA-transduced cells were sacrificed at 15 weeks after transplantation, without HIV challenge. Splenocytes and bone marrow cells were collected and tested for GFP expression, as a marker of shPromA. Cells were also checked for the expression of CD45 (hematopoietic cells), CD3 (T cells), CD4, CD8, CD19 (B cells), CD56 (subset of NK cells), CD123 (subset of dendritic cells), CD33 (myeloid cells), CD11c, CD66b (granulocytes), and CD14 (monocytes). There was no obvious skew in GFP expression among all phenotypically unique cells tested (Fig. 1B and D).

### HIV-1_JRFL_ challenge of NOJ mice transplanted with shRNA PromA-expressing CD34^+^ cells

The two mice groups (PromA and CD34) were challenged with HIV-1_JRFL_ at 14–16 weeks after transplantation (Fig. 1E). Peripheral blood was collected at weeks 1 and 2 post challenge. Mice were sacrificed at week 2, and splenocytes and bone marrow cells (from the left femur) were collected for further analysis (Fig. 1E). The frequencies of human CD45^+^ cells and their subsets were analyzed using PBMCs (weeks 1 and 2), splenocytes (week 2), and bone marrow cells (week 2) (Fig. S4, Table 1). In PBMCs, the percentage of human CD45^+^ cells was significantly lower in the PromA group than in the CD34 group (Fig. S4A). There was also a significant difference in counts of peripheral blood human CD45^+^ cells at week 1 (Fig. S4B). However, there was no significant difference in blood CD45^+^ cell counts at week 2 between the two groups (Fig. S4B). Moreover, the CD45^+^ cell counts in the spleen and bone marrow were much higher in the PromA group than that in the CD34 group (Table 1). The reasons for these discrepancies remain unclear.

**Table 1.**
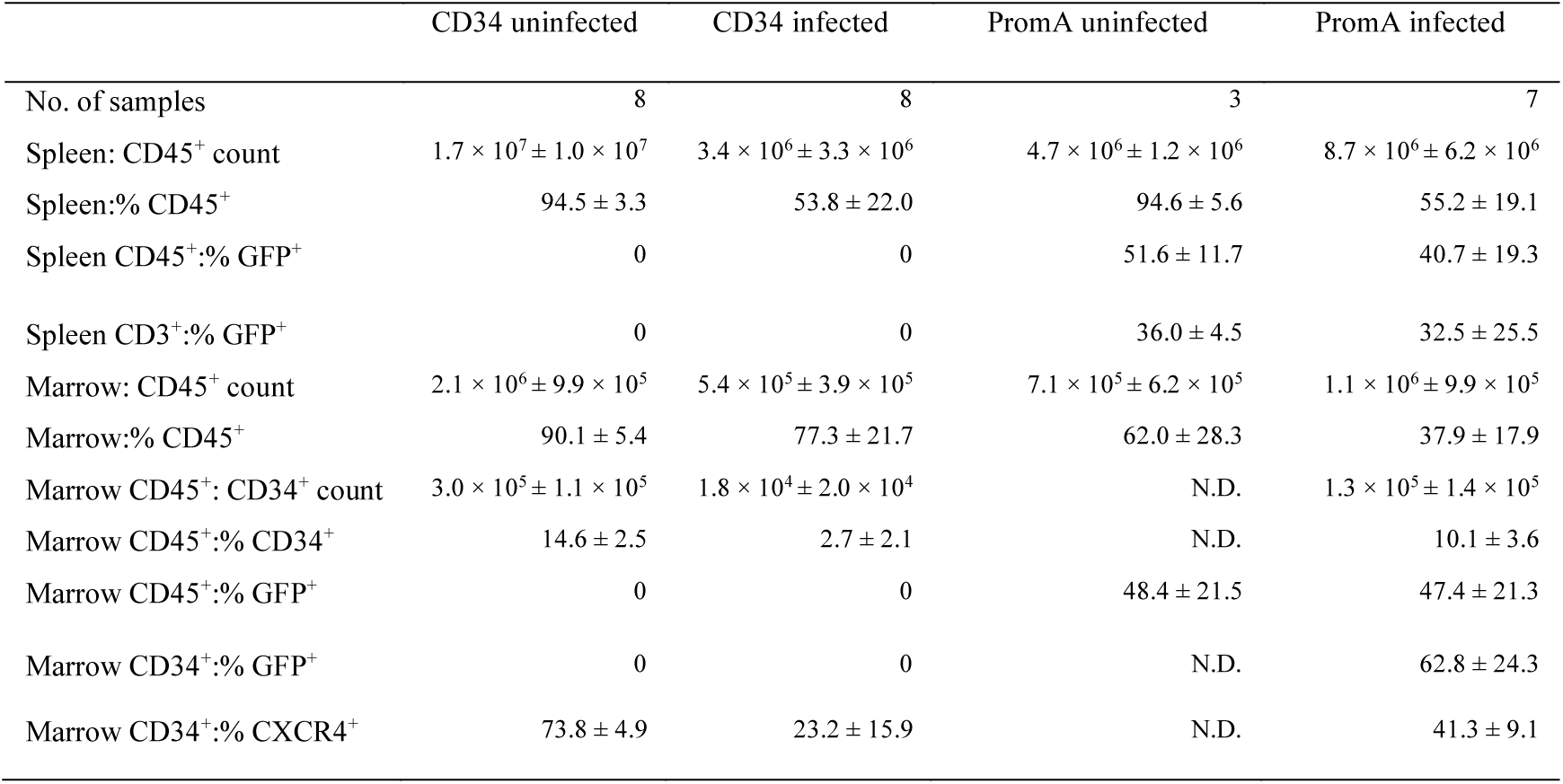
Comparison of spleen/bone marrow cells among the animal groups untransduced (CD34) uninfected, untransduced (CD34) infected, PromA-transduced uninfected, and PromA-transduced infected. Values are presented as mean ± standard deviation. Bone marrow cells were collected from the left femur of each animal. Samples were collected at week 15–18 post transplantation or at week 2 post challenge for HIV-infected animals (Fig. 3). N.D., not determined.

### CD4^+^CCR5^+^ T cell dynamics after HIV-1_JRFL_ challenge of humanized NOJ mice

Intracellular HIV Gag p24 expression levels were measured in PBMCs (weeks 1 and 2), splenocytes (week 2), and bone marrow cells (week 2) after HIV challenge. There were significant differences in HIV p24^+^ frequencies in CD3^+^CD8^‒^ T cells at weeks 1 and 2, indicating the impact of PromA on suppression of HIV replication (Fig. 2A and B, Fig. S5). Cell-associated HIV DNA/RNA copy numbers in splenocytes (week 2) relative to human GAPDH were measured by quantitative PCR/RT-PCR. Compared with those in the CD34 group, the average relative RNA copy number was low, whereas the relative DNA copy numbers were significantly high in the PromA group, (Fig. 2C left and middle). The ratios of HIV RNA to DNA levels were calculated. The average RNA/DNA ratio in the PromA group was >10-fold lower than in the CD34 group (Fig. 2C right), showing suppression of HIV RNA, including transcripts, compared with DNA. There was no mutation in the PromA region of the HIV RNA (data not shown).

**Fig 2.**
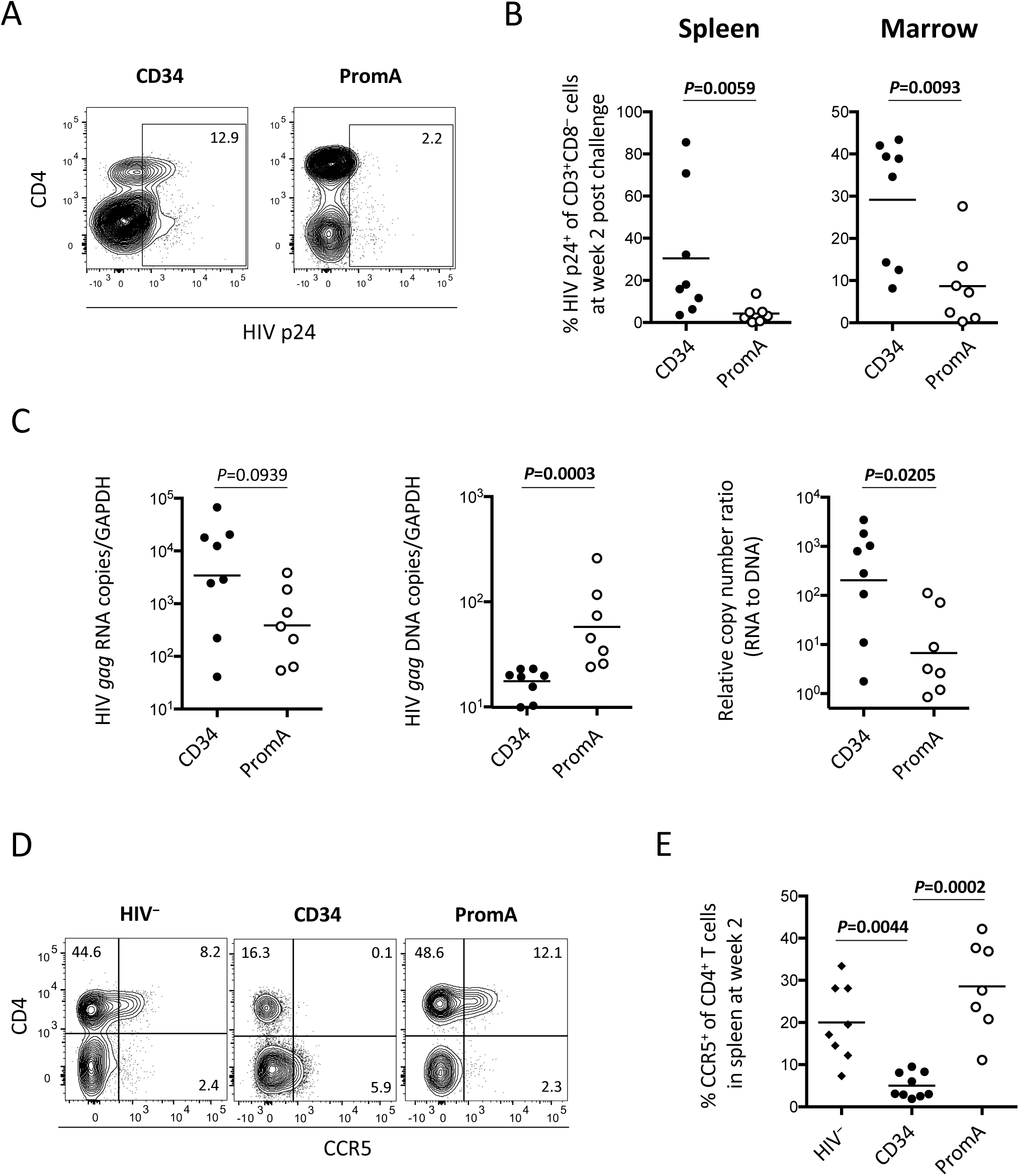
PromA suppressed viral replication and protected CD4^+^ T cells from depletion in NOJ mice. (A) Representative plots showing suppression of intracellular HIV p24 expression in PromA-transduced mice. (B) Intracellular HIV p24^+^ frequencies in CD3^+^CD8^-^T cells were compared between the CD34 and PromA groups. The analysis was done using splenocytes (left) and bone marrow cells (right) obtained at week 2 post challenge. (C) Cell-associated RNA (left) and DNA (middle) copies were normalized to GAPDH. Relative RNA copies per DNA were calculated from these data (right). Statistical analysis was performed using the Mann–Whitney U-test. (D) Representative plots showing preserved CD4^+^CCR5^+^ cells in the PromA-transduced mice compared with the control mice. (E) CCR5^+^ frequencies of CD4^+^ T cells collected from the spleen at week 2 post challenge. Statistical analysis was performed by the multiple comparison test with Dunn’s method.

There was a marked decrease in the numbers of CD4^+^CCR5^+^ splenocytes in the CD34 group at week 2, compared with the uninfected control group (Fig. 2D and E). Conversely, splenic CD4^+^CCR5^+^ T-cell frequencies were preserved in the PromA group, supporting the protective effect of PromA against challenge with the CCR5-tropic strain of HIV-1 (Fig. 2D and E). Analysis of PBMCs sampled at week 2 post challenge showed similar results (Fig. S6).

### Bone marrow CD34^+^ cell dynamics in humanized NOJ mice after challenge with CCR5-tropic HIV-1_JRFL_

To better clarify the protective effect of PromA on human hematopoietic cells in HIV infection, bone marrow CD34^+^ cells were analyzed (Fig. 3). At week 2 post challenge, the mean percentage of bone marrow CD34^+^ cells was severely decreased in the CD34 group, compared with the uninfected control group (2.7% vs. 14.6%, respectively) (Fig. 3A–D). Further analysis revealed preferential depletion of CD34^+^CXCR4^+^ cells, as shown by the drop in the mean percentage of CXCR4^+^ cells in the CD34^+^ cell population from 73.8% (HIV^-^) to 23.2% (CD34) (Fig 3A–D). The mean percentage of CD34^+^ cells in the PromA group was significantly greater than in the CD34 group (10.1% vs. 2.7%, respectively, Fig. 3D). This finding indicated that shPromA was associated with partial protection of bone marrow CD34^+^ cells, including the CD34^+^CXCR4^+^ cell population (Fig 3A–D). The expression levels of shPromA (GFP) in the bone marrow cells of mice in the PromA group were further analyzed. Interestingly, a greater percentage of CD34^+^ cells were positive for GFP compared with CD45^+^ cells (Fig. 3F), implicating a possible role of shPromA in the protection of CD34^+^ cells from HIV infection.

### In vitro-generated PromA-expressing macrophages were resistant to HIV-1_JRFL_ replication

In parallel with the HIV challenge experiment using the humanized mice, the effect of shPromA on suppression of HIV replication was tested with macrophages generated *in vitro* from cord-derived CD34^+^ cells (Fig. 4A). The average percentage of GFP^+^ cells in CD11b^+^ cell populations was 42.6%, and most CD11b^+^ cells were CD14^+^/CD16^+^/CD4^+^/CCR5^+^ (Fig. 4B). Five days after infection of these macrophages with HIV-1_JRFL_, viral replication levels, measured as intracellular HIV p24^+^ percentages, were compared among untransduced, shPromA-M2-transduced, and shPromA-transduced macrophages. As shown in Fig 4C and D, HIV replication was significantly suppressed in shPromA-transduced macrophages, compared with autologous control samples. Similar results were obtained with PM1-CCR5 cells transduced with shPromA or shPromA-M2 and infected with HIV-1_JRFL_ (Fig. S7).

**Fig 4.**
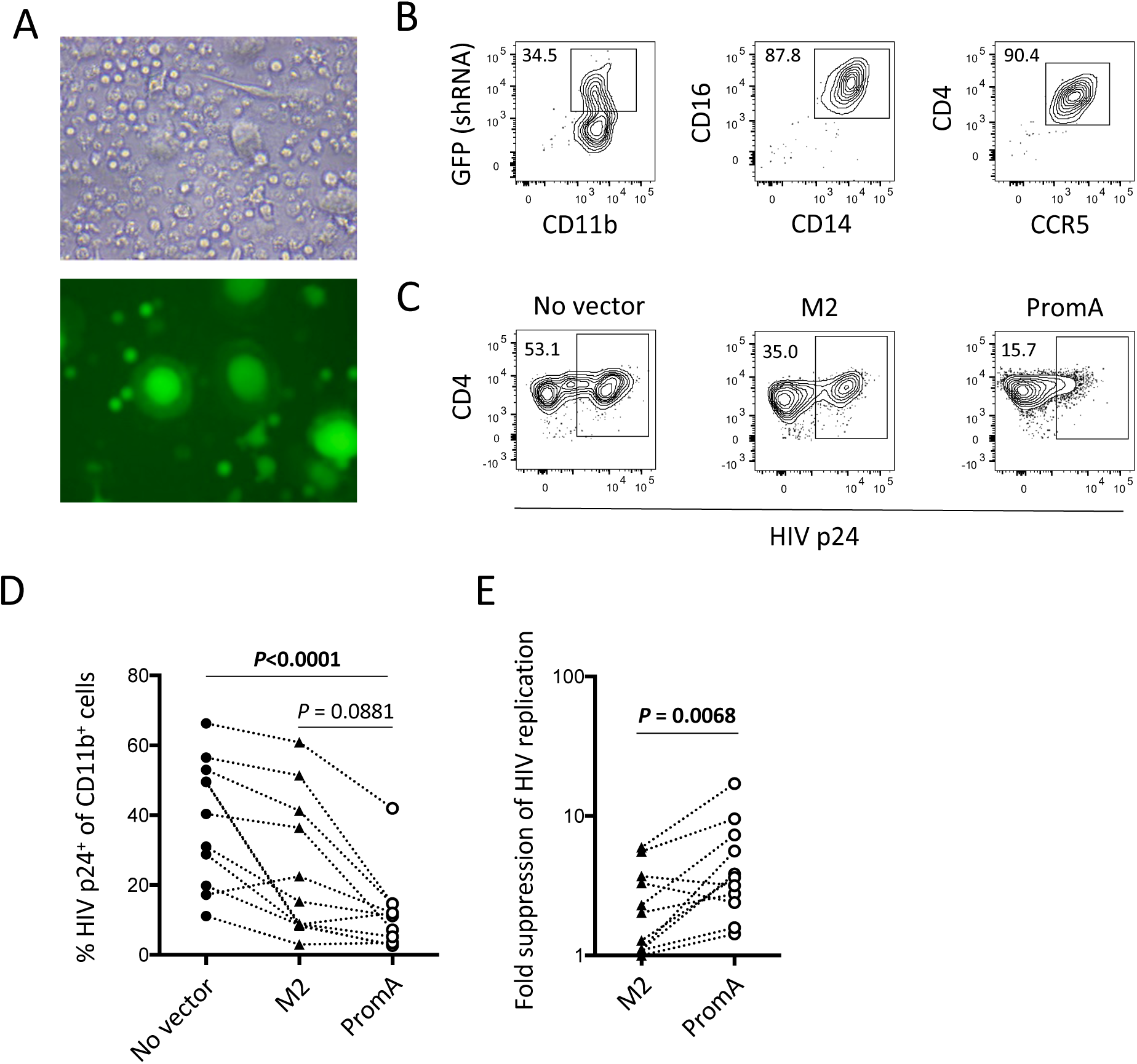
Freshly isolated CD34^+^ cells were transduced with shPromA and differentiated to macrophages *in vitro*. Cells were then infected with HIV-1_JRFL_. HIV replication levels were compared to autologous mock-transduced or shPromA-M2-transduced macrophages on day 5 post infection. (A) PromA-expressing (GFP^+^) macrophages with the typical fried-egg shapes were observed by fluorescence microscopy. (B) Representative plots showing expression patterns of GFP (PromA)/CD11b (left) in live cells as well as expression patterns of the CD14/CD16 (middle) and CD4/CCR5 (right) subets in the population of CD11b^+^ cells. (C) Representative plots showing intracellular HIV p24^+^ frequencies in mock-transduced, shPromA M2-transduced, and shPromA-transduced macrophages on day 5 post infection. (D) HIV replication was significantly lower in the shPromA-transduced samples than in mock-transduced samples. Nonparametric multiple comparison analysis was performed with the Dunn’s method. (E) Fold suppression of HIV replication relative to mock-transduced samples was calculated for PromA M2-transduced and shPromA-transduced samples. The shPromA-transduced macrophages showed a significantly higher suppressive capacity. Comparisons were performed with the Wilcoxon matched-pairs signed rank test or Dunn’s method of multiple comparison analysis.

## Discussion

One of the challenges with standard HIV therapies is the lack of an effective method to repair the compromised immune system during treatment initiation, regardless of AIDS manifestation [20]. Gene therapy with hematopoietic stem/progenitor cells (HSPCs) has been of great interest because this can possibly recover the loss of the naïve/memory T-cell repertoire, which then would enable re-immunization of the individual against various pathogens, including HIV [21]. Thus far, treatments using knockout or knockdown of CCR5, a coreceptor used by strains of HIV-1 that preferentially infect memory CD4+ T cells, are the most successful [22]. In addition, other potential host and non-host targets to render the host cells more resistant to HIV replication have been investigated [23]. The PromA region in the HIV genome has proved to be among the most safe and powerful non-host targets [24].

In the present study, newborn NOJ mice were transplanted with lentivirally transduced CD34^+^ cells expressing shPromA, which resulted in shPromA-expressing CD34^+^ cell engraftment and their differentiation to most major subtypes of myeloid and lymphoid cells *in vivo*, including CD4^+^/CD8^+^ T cells, CD19^+^ B cells, CD56^+^ NK cells, and CD11c^+^/CD14^+^/CD66b^+^/CD123^+^ myeloid cells (Fig. 1B–D, Table 1). These findings suggested that the off-target effects associated with PromA expression did not interfere with hematopoietic cell differentiation *in vivo* [13]. The method tested in this study is thus promising and also in line with previous reports showing that retroviral transduction does not alter immune function [25].

The results of the HIV-1_JRFL_ challenge experiment showed that the HIV-infected mice were depleted of CD4^+^CCR5^+^ cells and bone marrow CD34^+^ cells. Interestingly, depletion of the latter occurred in a CXCR4-associated manner (Fig. 3A–D), which may implicate increased CD34^+^ cell turnover mediated by the CXCR4/SDF-1 signaling pathway, compensating for the increased CD4^+^ cell turnover following HIV infection. Although CXCR4-tropic HIV-1 infects HSPCs more efficiently than CCR5-tropic HIV-1 [26], CCR5-tropic HIV-1 alone may be sufficient to cause bone marrow dysfunction associated with immunological nonresponsiveness and/or pancytopenia [3, 4, 27]. In addition, in the CD34 group, bone marrow CD34^+^ cell depletion was more profound than spleen/bone marrow CD45^+^ cell depletion (Table 1). This may suggest early involvement of bone marrow dysfunction in disease progression, such as impaired CD4^+^ T-cell production, rather than being only a determinant of the immune reconstitution rates in the chronic phase.

Conversely, HIV replication was suppressed (Fig. 2A-C) and CD4^+^CCR5^+^ T cells were preserved (Fig. 2D and E, Fig. S6) in mice engrafted with shPromA-transduced cells; this is relevant to the previous *in vivo* results using adult NOJ mice transplanted with shPromA-transduced PBMCs [13]. Mice in the PromA group also showed partial preservation of bone marrow CD34^+^ cells (Fig. 3), which implies that the anti-HIV gene therapy of CD34^+^ HSPCs may be more advantageous than equivalent gene therapy of PBMCs [20, 28]. Multiple types of hematopoietic progenitor cells may harbor CCR5-or CXCR4-tropic viruses for extended periods [29]; therefore, the expression of an anti-HIV modality in these cell types may be highly effective to protect the hematopoietic system of HIV-infected patients.

Limitations of this study include the lack of an empty lentivirus vector for the patented shPromA lentivirus vector. Instead, mice transplanted with unmanipulated CD34^+^ cells were included as a control group. The number of animals was too limited to create another group of mice transplanted with CD34^+^ cells expressing shPromA-M2 with a three-base mismatch, compared with shPromA. To partly compensate for this limitation, *in vitro* viral suppression assays with macrophages (Fig. 4) were conducted using cells either untransduced or transduced with shPromA-M2 as controls. Another limitation is that the HIV challenge was conducted as a 2-week acute infection experiment with injection of a high dose of HIV-1_JRFL_ to each animal. A chronic infection study with an injection of a low dose of HIV-1_JRFL_ may be useful to determine whether PromA is able to support the preservation and function of bone marrow CD34^+^ cells for prolonged periods [30].

In summary, the present *in vivo* study of humanized mice demonstrated that infection of CCR5-tropic HIV-1 alone resulted in depletion of bone marrow CD34^+^ stem/progenitor cells in a CXCR4-associated manner. Gene therapy of HSPCs with shPromA targeting a non-host, non-coding gene sequence may be beneficial to preserve the hematopoietic potential of the bone marrow [16], which would be a substantial step toward the clinical application of TGS (e.g., combination of PromA, targeting of CCR5, and antiretroviral therapies).

## Acknowledgments

I thank Drs. Kazuo Matsui and Shoichi Kawakami of Fukuda Hospital, Kumamoto, Japan, for their assistance with cord blood sampling. I thank Ms. Michelle Millington, Ms. Maureen Boyd, and Dr. Geoff Symonds for providing lentiviral vector constructs and the transduction protocol. I thank Prof. Anthony Kelleher, Dr. Kazuo Suzuki, and Ms. Katherine Marks of the Kirby Institute, UNSW, Australia, for their support and help with preparation of the shRNA-expressing lentivirus vectors. I thank Prof. Seiji Okada of the Center for AIDS Research, Kumamoto University, and his staff for their technical assistance with the animal experiments. I thank Enago (www.enago.jp) for the English language review. This work was supported by grants from the National Health and Medical Research Council, Australia (Project grant), the Japan Agency for Medical Research and Development (Research program on HIV/AIDS, no. 16fk0410108h0001), and the Ministry of Education, Culture, Sports, Science and Technology, Japan (Grants-in-Aid for Science Research, no. 25114711).

---

**Fig. S1.**
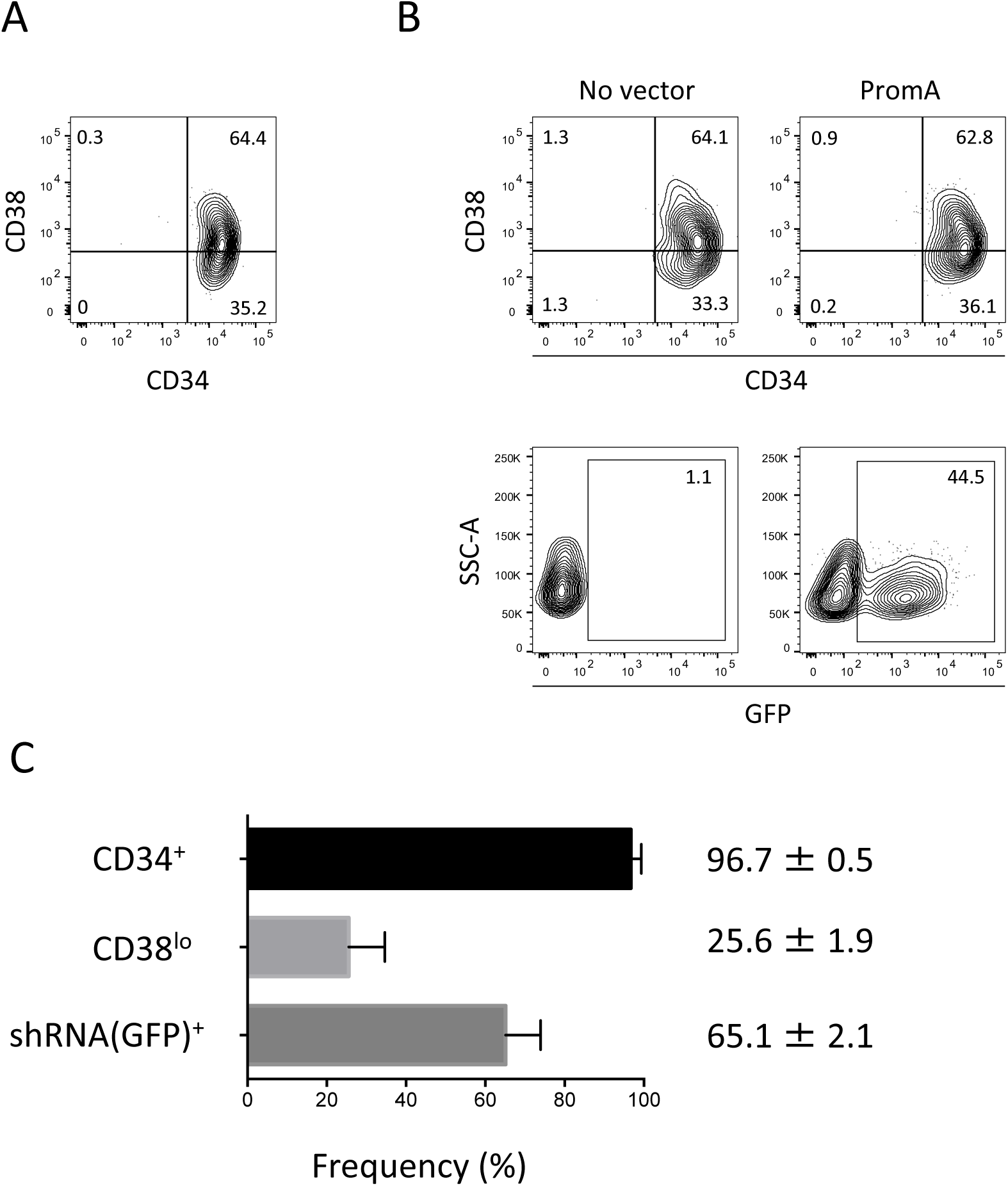
Fresh cord-derived CD34^+^ cells were isolated and transduced with a lentivirus vector expressing the shRNA PromA adapted to HIV-1_JRFL_ (shPromA_JRFL_ or simply shPromA). (A) A representative plot showing the CD34/CD38 expression pattern in freshly isolated CD34^+^ cells. (B) Representative plots showing the lentiviral transduction efficiency measured by GFP expression. (D) Summary of the CD34^+^ cell isolation results (upper and middle bars) and the efficiency of the following lentiviral transduction (lower bar).

**Fig. S2.**
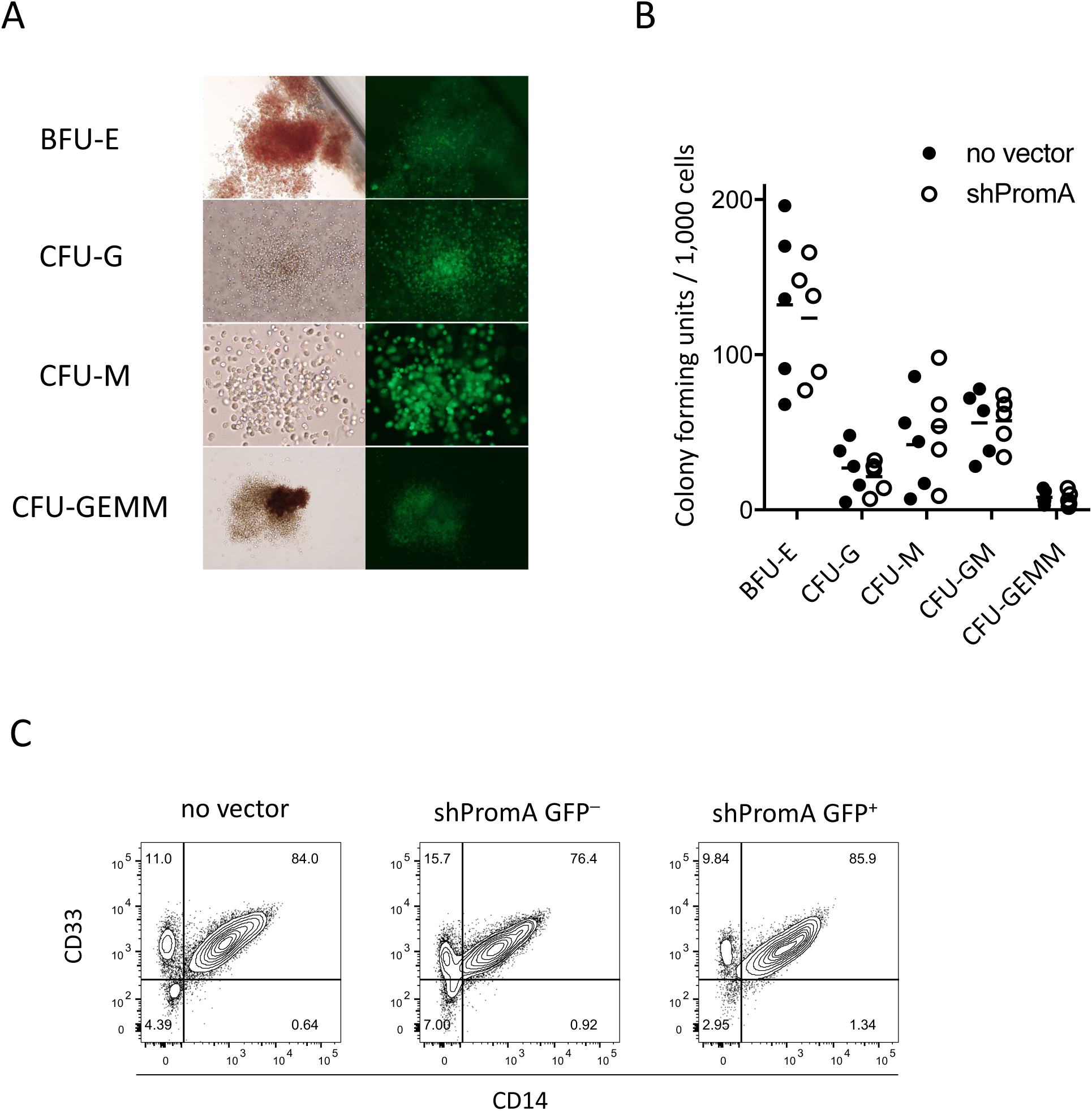
PromA-transduced CD34^+^ cells showed equivalent colony forming capability to mock-transduced CD34^+^ cells. In vitro differentiation of shPromA-transduced CD34^+^ cells to erythrocytes, granulocytes, or monocytes were assessed in vitro by a colony formation assay. Cells were diluted to 500 /mL in MethoCult Optimum methylcellulose media (STEMCELL Technologies, VIC, Australia) containing recombinant human cytokines including SCF, granulocyte macrophage-colony stimulating factor (GM-CSF), granulocyte-colony stimulating factor (G-CSF), interleukin 3 (IL-3), and erythropoietin. Cells were seeded in 6-well plate (Corning, VIC, Australia) using a syringe and BD blunt plastic cannula (BD Biosciences, NSW, Australia). Colony-forming units per 1,000 CD34^+^ cells were counted after 14 days of culture. (A) Different types of GFP^+^ colonies were found. (B) PromA-transduced CD34^+^ cells showed no significant difference to mock-transduced cells in colony forming units of all the different colony types tested. (C) Colony forming cells were collected together for flow cytometric analysis. CD14/CD33 expression levels were compared between mock-transduced, shPromA-transduced GFP^-^, and PromA-tranduced GFP^+^ cells.

**Fig. S3.**
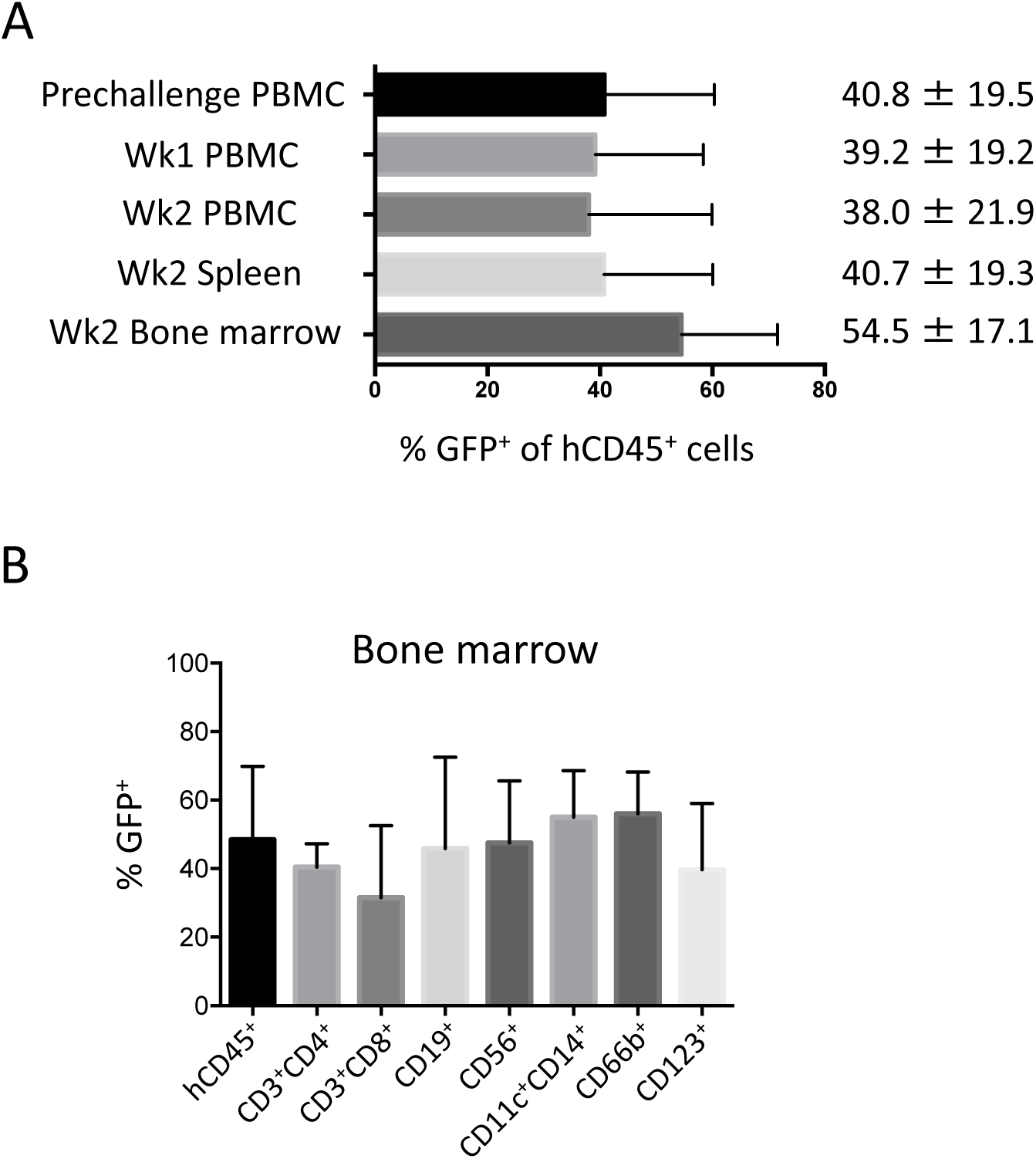
Engraftment of human cells in three transplanted NOJ mice was tested 15 weeks after transplantation. (A) PromA-expressing frequencies in human CD45^+^ cells, expressed as GFP^+^ frequencies, in group PromA are shown. These were tested in mouse PBMC (prechallenge, week 1, week 2), splenocytes (week 2) and bone marrow cells (week 2) (B) The mean GFP^+^ percentages in the different subsets of cells in bone marrow.

**Fig. S4.**
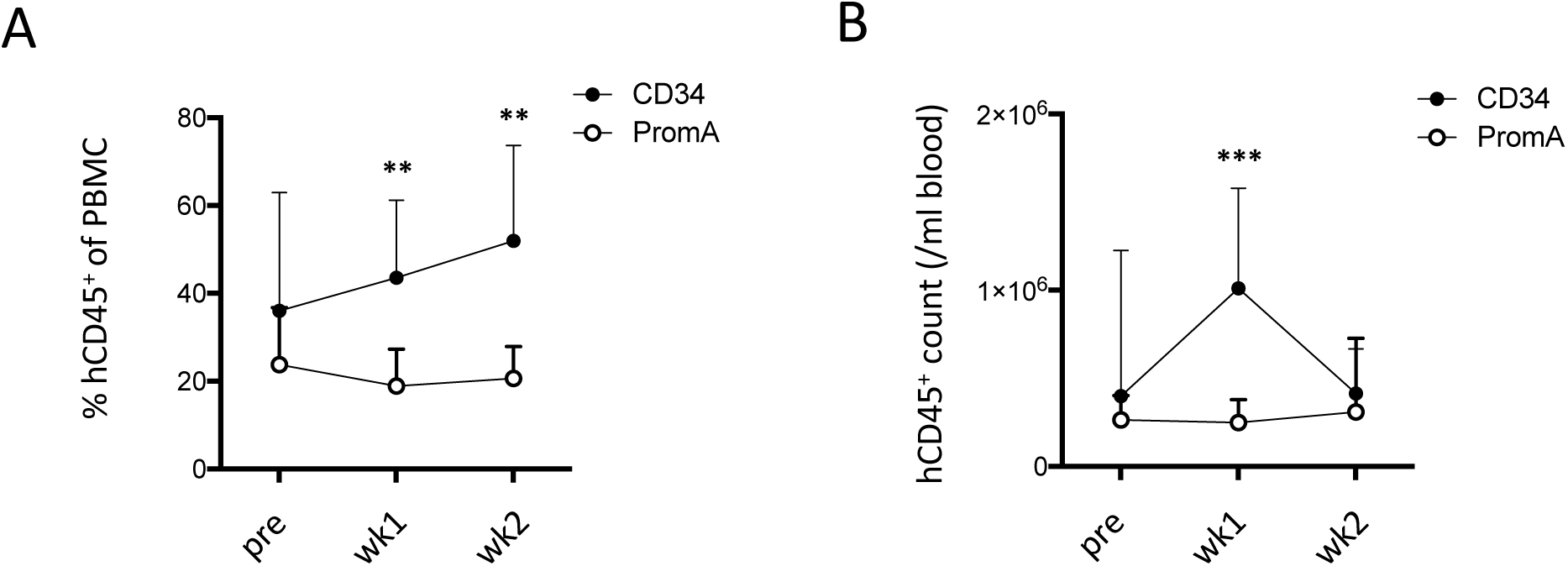
Blood CD45^+^ frequencies were analyzed after HIV-1 infection of humanized mice. (A) Human CD45^+^ percentages in PBMC at prechallenge, week 1, and week 2. (B) Human CD45^+^ cell counts in blood (/ml) at prechallenge, week 1, and week 2. Comparison was done by the Mann-Whitney test. **: *P*<0.01, ***: *P*<0.001.

**Fig. S5.**
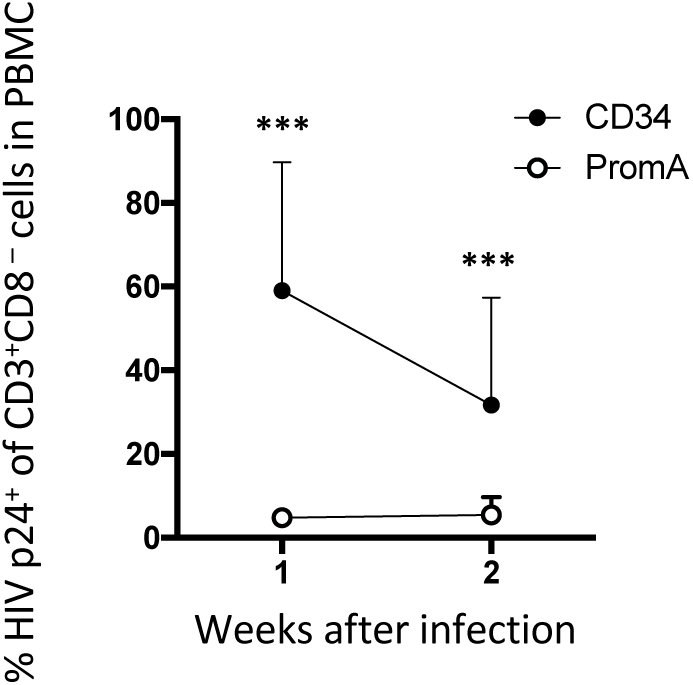
Intracellular HIV p24^+^ frequencies in CD3^+^CD8^-^ T cells were compared between group CD34 and PromA of HIV-infected humanized mice. The analysis was done using PBMC obtained at weeks 1 and 2 post challenge. Comparison was done by the Mann-Whitney test. ***: *P*<0.001.

**Fig. S6.**
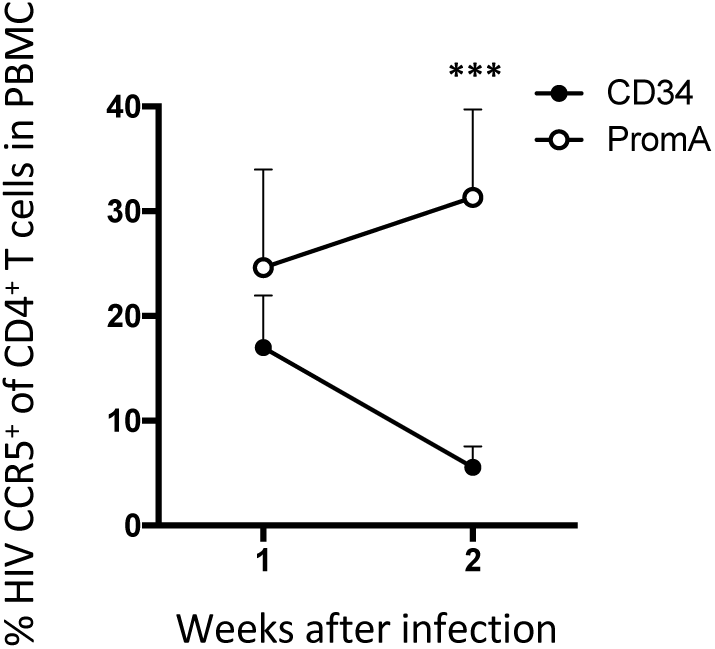
CCR5^+^ T-cell frequencies of CD4^+^ T cells in PBMCs were analyzed at weeks 1 and 2 post HIV challenge of humanized mice. Comparison was done by the Mann-Whitney test. ***: *P*<0.001.

**Fig. S7.**
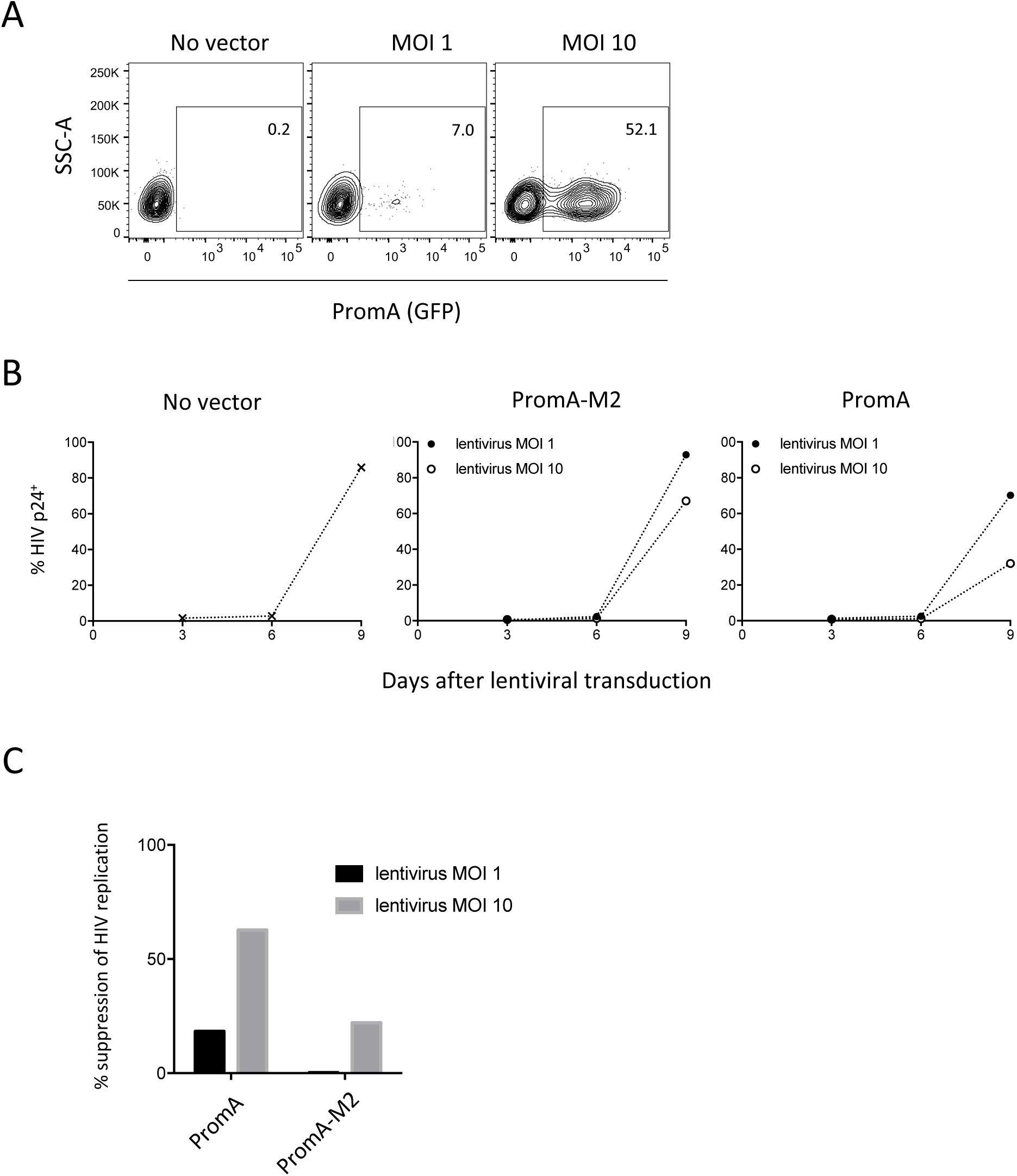
The efficacy of shPromA to suppress HIV infection was tested in vitro using the PM1-CCR5 cells^1^. Cells were infected with HIV-1_JRFL_ at an MOI of 0.01. Twenty-four hours later, cells were seeded in RetroNectin-coated 48-well plate at the concentration of 5,000 cells/well and transduced with the lentivirus expressing shPromA at an MOI of 1 or 10. Cells were analyzed at days 3, 6, and 9 post-transduction for the intracellular expression of HIV p24^+^. (A) Transduction rates as determined by the expression of GFP analyzed by flow cytometry. The lentiviral MOIs were indicated. (B) Intracellular HIV p24^+^ percentages monitored at days 3, 6, and 9 post-transduction. (C) Percentages of suppression of HIV replication relative to the untransduced control at day 9 post-transduction were calculated by the data shown in B.

## Footnotes

1 Yusa, K., Maeda, Y, Fujioka, A., Monde, K., Harada, S., 2005. Isolation of TAK-779-resistant HIV-1 from an R5 HIV-1 GP120 V3 loop library. J Biol Chem 280, 30083-30090.

